## Supplementary Figures for "Host environment shapes filarial parasite fitness and *Wolbachia* endosymbionts dynamics"

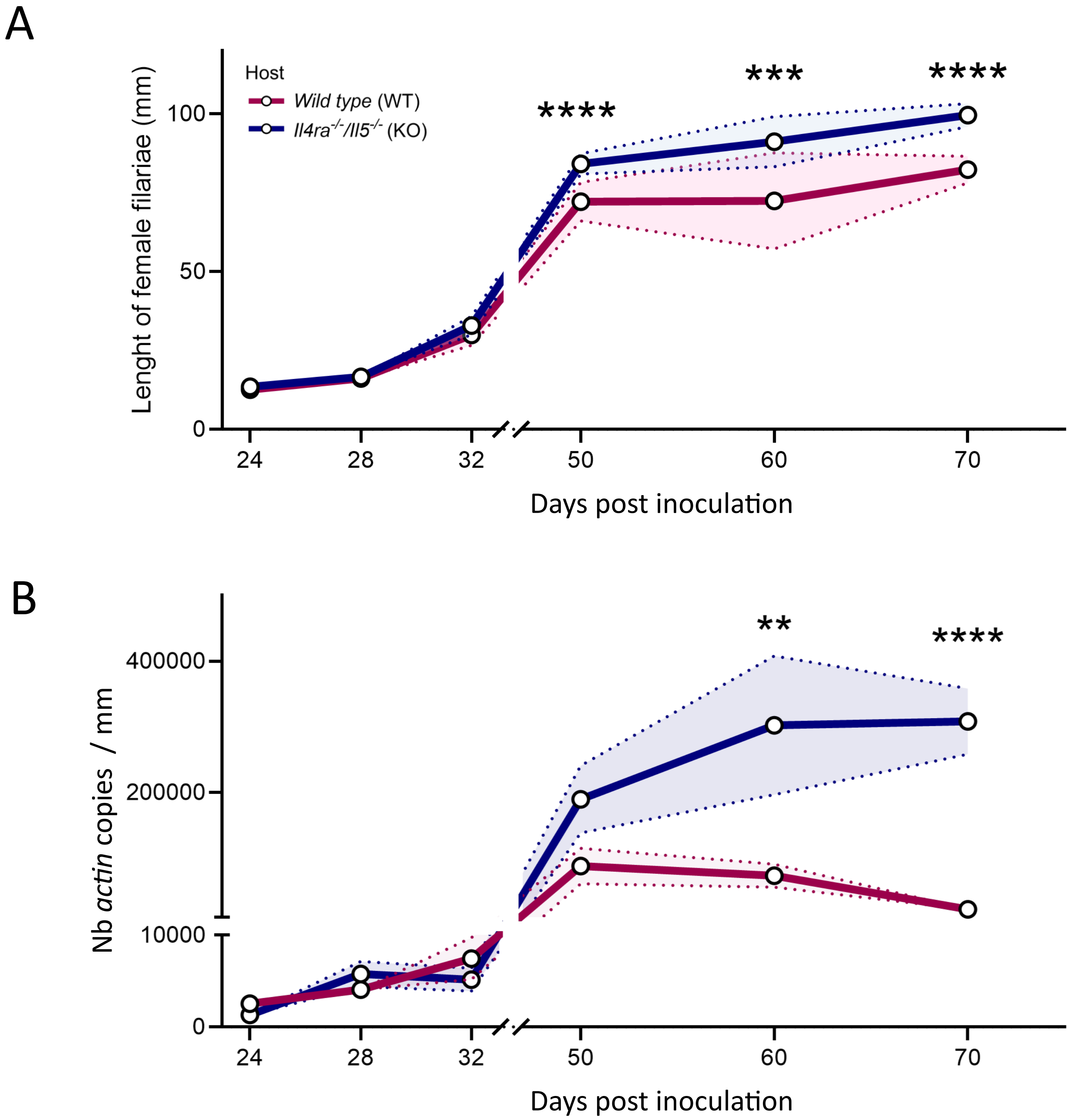


**Supplementary Figure 1: Growth and *actin* gene expression of female *Litomosoides sigmodontis* in wild-type and type 2-deficient mice.** Type 2-competent wild-type (WT) and type 2-deficient (*Il4rα$^{-/-}$/Il5$^{-/-}$, KO*) mice were inoculated with 40 infective larvae (L3) of the filaria *L. sigmodontis*. Parasites were harvested and measured at various time points before and after the fourth molt (~30 dpi), and *Wolbachia's* gene *ftsZ* and filarial *actin* were evaluated by qPCR in female filariae. **(A)** Measurements of worm length in millimeters (mm) from 24 to 70 days post-infection (dpi) in wild-type and type 2-deficient (*Il4rα$^{-/-}$/Il5$^{-/-}$*) mice. Results are expressed as the mean ± SD of n= 20-40 filariae per group (24-50 dpi), and n = 8 filariae from wild-type and 28 filariae from *Il4rα$^{-/-}$/Il5$^{-/-}$* hosts (60 dpi) . Two-way ANOVAs followed by Bonferroni's multiple comparisons tests were performed; ***p < 0.001, ****p < 0.0001 indicate significant difference between filariae from wild-type and *Il4rα$^{-/-}$/Il5$^{-/-}$* hosts. **(B)** Quantification of *actin* gene expression, normalized to worm length, over the same period. These measurements provide a baseline for the relative quantification of *Wolbachia* density shown in Figure 1, accounting for changes in worm size that could influence the interpretation of bacterial load. Results are expressed as the mean ± SD of n = 4-6 filariae per group (24-60 dpi), n = 11-15 filariae per group (70 dpi). Two-way ANOVAs followed by Bonferroni's multiple comparisons tests were performed; **p < 0.01, ***p < 0.001 indicate significant difference between filariae from wild-type and *Il4rα$^{-/-}$/Il5$^{-/-}$* hosts.

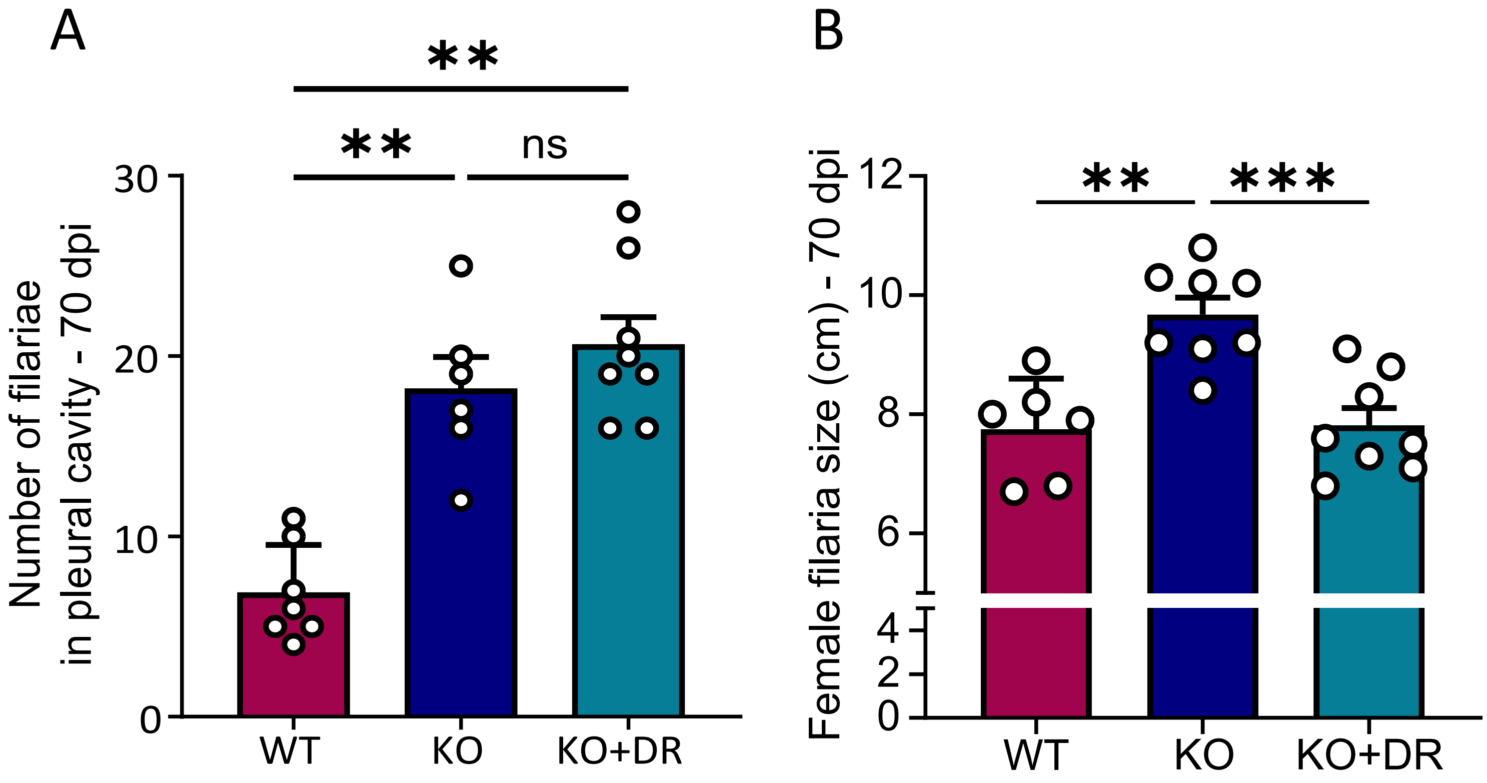


**Supplementary Figure 2: Effect of host immune background and *Wolbachia* depletion on filarial size and pleural cavity burden at 70 dpi. (A)** Number of filariae recovered in the pleural cavity of WT, KO, and KO+DR mice at 70 dpi. Results are expressed as mean ± SEM, with individual data points representing each mouse (n = 6–8 mice per group). One-way ANOVA followed by Tukey's multiple comparison test were performed. **$p < 0.01$, ns: not significant.

**(B)** Female filariae size (cm) from WT, KO, and KO+DR mice at 70 dpi. Results are expressed as mean ± SEM, with individual data points representing each measured worm (n = 6–8 worms per group). One-way ANOVA followed by Tukey's multiple comparison test were performed. **$p < 0.01$, ***$p < 0.001$.

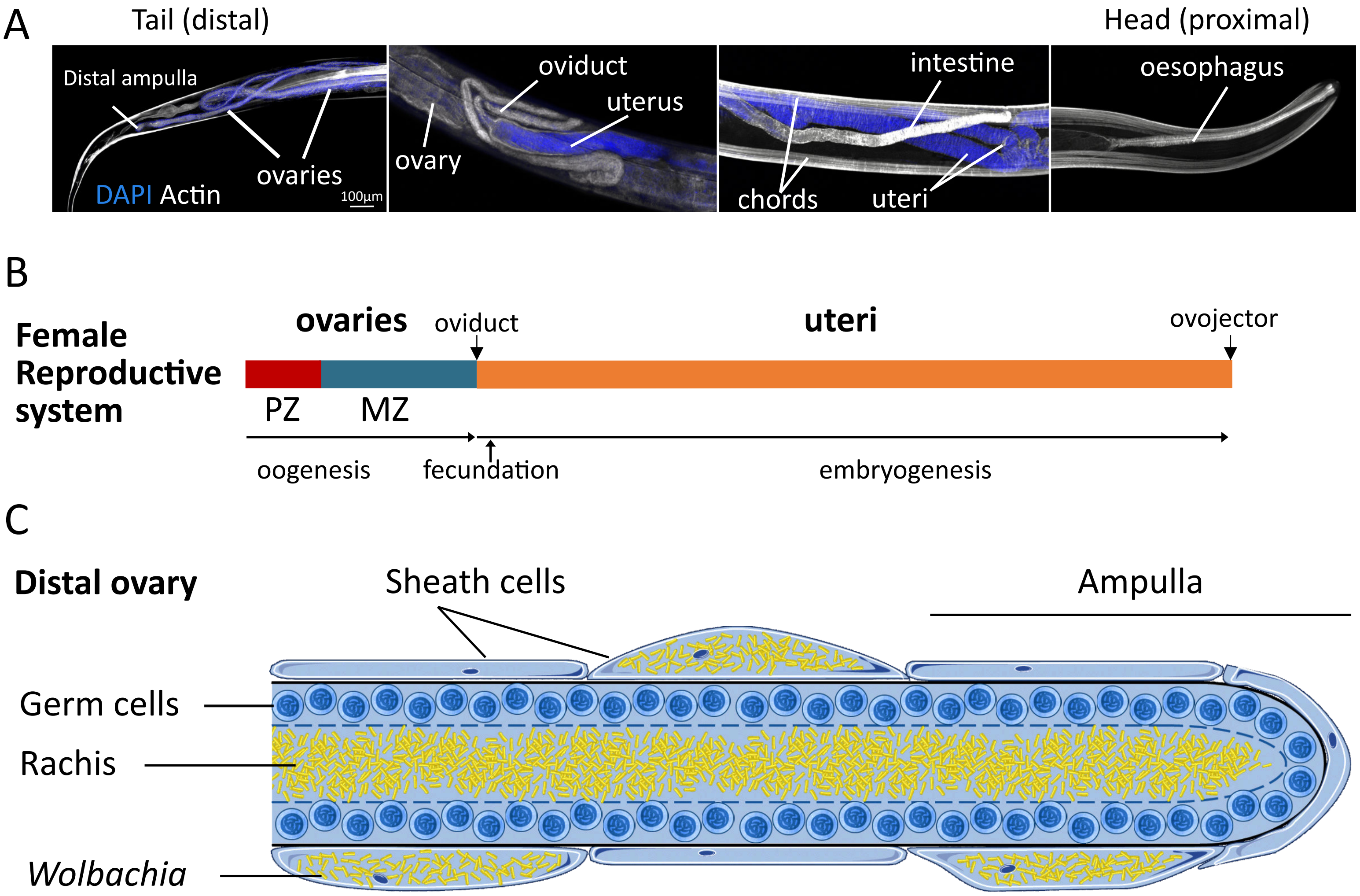


**Supplementary Figure 3. Anatomical organization of the female reproductive system and *Wolbachia* localization in *Litomosoides sigmodontis*. (A)** Fluorescent microscopy images of an adult female filaria highlighting key anatomical regions. The distal ampulla and ovaries are situated at the posterior end, transitioning through the oviduct into the uteri, which extend along the body axis. The proximal uteri, near to the ovojector, contain mature microfilariae. Staining highlights nuclei (DAPI, blue) and actin filaments (gray). **(B)** Schematic representation of the female reproductive system. The ovary includes a proliferative zone (PZ, red), where germ cells divide, and a meiotic zone (MZ, blue), where meiosis occurs. The uteri (orange) extend anteriorly and terminate at the ovojector, where mature microfilariae are expelled. **(C)** Diagram of the distal ovary, showing the rachis (central cytoplasmic core) surrounded by germ cells. *Wolbachia* (yellow) are distributed along the rachis and within some sheath cells. The ampulla marks the distal tip of the ovary and the starting point of oogenesis. In somatic tissues, *Wolbachia* also reside in the lateral hypodermal chords (see Figure 2E).

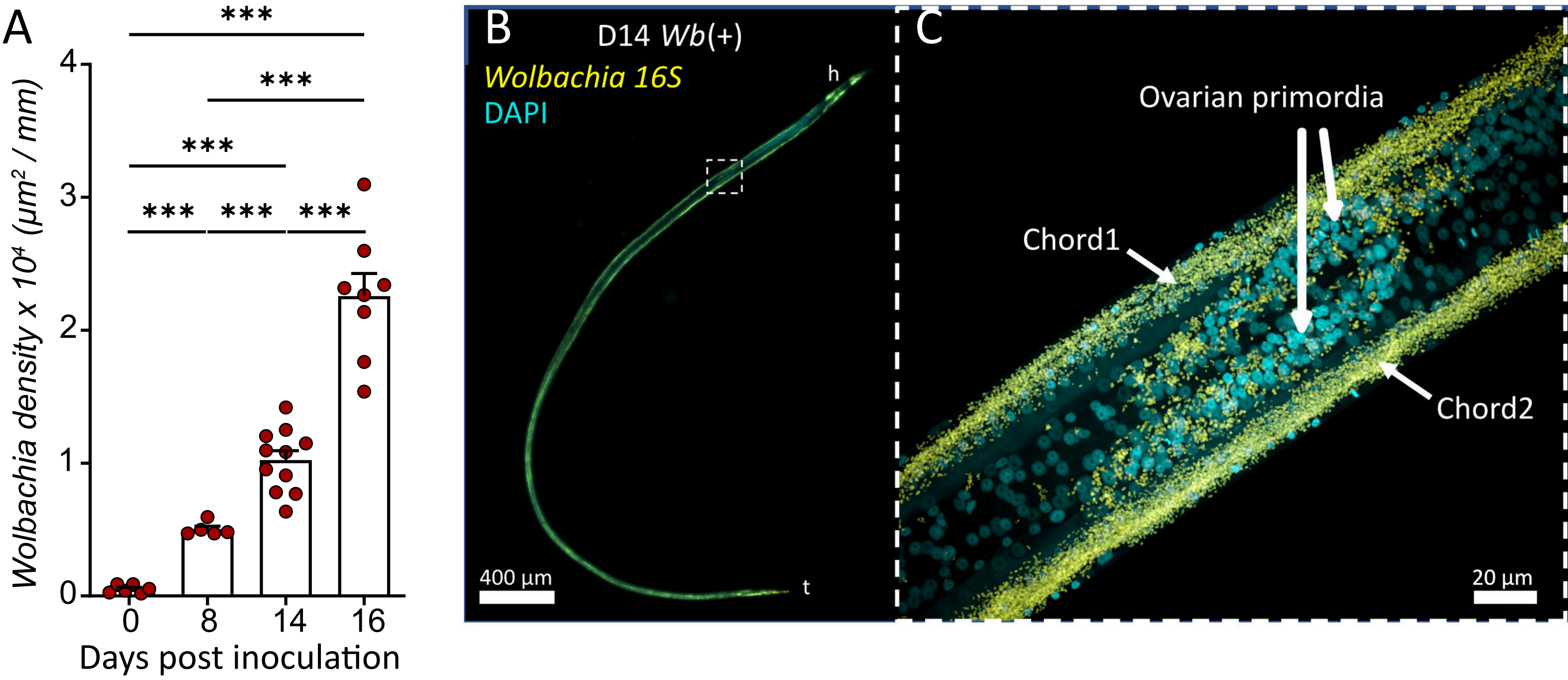


**Supplementary Figure 4. Colonization of the ovarian primordia by *Wolbachia* in L4 female filariae. (A)** Quantification of *Wolbachia* density (µm²/mm) in *Litomosoides sigmodontis Wb*(+) larvae at different days post-inoculation, based on fluorescence microscopy images of entire larvae. Brown-Forsythe ANOVA test followed by a Dunnett's T3 multiple comparisons post-hoc test (****p* < 0.001). **(B)** Confocal fluorescence image of a *Wb*(+) L4 female *L. sigmodontis* at day 14 post-infection (D14). The worm was stained for DNA (DAPI, cyan) and *Wolbachia* (16S ribosomal RNA, yellow). The anterior-to-posterior orientation is indicated (h = head; t = tail). **(C)** Magnified region showing the lateral hypodermal chords and ovarian primordia in the boxed area from (B). *Wolbachia* are visible in the lateral chords (Chord1 and Chord2) and are invading the ovarian primordia.
